# Transcriptional maintenance of cortical somatostatin interneuron subtype identity during migration

**DOI:** 10.1101/593285

**Authors:** Hermany Munguba, Kasra Nikouei, Hannah Hochgerner, Polina Oberst, Alexandra Kouznetsova, Jesper Ryge, Renata Batista-Brito, Ana Belén Munoz-Manchado, Jennie Close, Sten Linnarsson, Jens Hjerling-Leffler

## Abstract

Recent work suggests that cortical interneuron diversity arises from genetic mechanisms guided by the interplay of intrinsic developmental patterning and local extrinsic cues. Individual genetic programs underlying subtype identity are at least partly established in postmitotic neural precursors, prior to their tangential migration and integration in the cortical circuitry. Nevertheless, it is unclear how distinct interneuron identities are maintained during their migration and maturation. Sox6 is a transcription factor with an established role in MGE-derived interneuron maturation and positional identity. To determine its role in maintaining somatostatin (Sst)-expressing interneurons’ subtype identity, we conditionally removed Sox6 in migrating Sst interneurons and assessed the effects on their mature identity using single-cell RNA-sequencing (scRNAseq), *in situ* hybridization and electrophysiology. Sox6 removal prior to migration in Sst-expressing neurons reduced subtype diversity without affecting overall number of neurons. Seven out of nine Sst-expressing molecular subtypes were absent in the mature primary somatosensory cortex of Sox6-cKO mice, including the Chodl-Nos1-expressing type which has been shown to be specified at, or shortly after, cell cycle exit. The remaining Sst-expressing subtypes in the Sox6-cKO cortex comprised three molecular subtypes, Crh-C1ql3 and Hpse-Cbln4, and a third subtype that seemed to be a molecular hybrid of these subtypes. Moreover, Sox6-cKO cells still expressed genes enriched within the entire class of Sst-expressing neurons, such as *Sst, Lhx6, Satb1, Elfn1* and *Mafb*. Removal of Sox6 at P7, after cells have reached their final destination and begin integration into the network, did not disrupt Chodl-Nos1 marker expression. Our findings suggest that expression of Sox6 during the migratory phase of cortical interneurons is necessary for maintenance of Sst^+^ subtype identity, indicating that subtype maintenance during migration requires active transcriptional programs.

## Introduction

Nascent neurons undergo incremental developmental steps that direct and restrict their fate until they reach terminal differentiation. During this process transcription factors are known to be functionally linked to inducing and maintaining the molecular blueprint acquired at previous developmental stages, by regulating gene expression, recruiting complementary transcriptional factors, and facilitating epigenetic modifications (Holmberg and Perlmann, 2012).

For example, the transcription factors network responsible for the differentiation of all 118 neuronal types present in the nematode *C. elegans* is well-characterized, ranging from transcription factors engaged in initial pan-neuronal specification through those which activate distinct neurotransmitter pathways (Hobert, 2016). As in invertebrates, mammalian neurons undergo a series of developmental steps before reaching their ultimate subtype identity, which consists of a distinct set of morphological, electrophysiological, molecular and connectivity features. In the mouse cortex alone, well over a hundred cell types have been identified (Tasic et al., 2018).

Instructive morphogens at the germinal zone induce a sequential cascade of transcription factors that shape neuronal identity. Initially, broad classes are established, e.g. glutamatergic *versus* GABAergic neurons or broad classes of GABAergic interneurons (derived from either the medial or caudal ganglionic eminences; MGE and CGE respectively). Thereafter, the process by which GABAergic cells from the same germinal zone acquire their discrete interneuronal types, e.g. MGE-derived parvalbumin-(Pvalb) *versus* MGE-derived somatostatin-(Sst) expressing GABAergic neurons is less clear (Hu et al., 2017; Inan et al., 2012). Yet another level of diversity exists, within which six to fifteen different Sst-expressing neuronal subtypes have been characterized according to morphological, functional and/or transcriptomic features (Munoz et al., 2017; Nigro et al., 2018; Tasic et al., 2018). This final level of subtype complexity was revealed only recently by high-resolution methods such as single-cell RNA-sequencing (scRNAseq), which made it possible to characterize the transcriptomic signatures of discrete cell types.

Recent work has demonstrated that newly postmitotic cortical Sst-expressing interneurons possess a unique transcriptomic signature beyond that expected for broad class identity and suggestive of their mature subtype identity (Mayer et al., 2018; Mi et al., 2018). This suggests that neuronal diversity is generated in the presence of local cues in the subventricular zone and is not completely dependent on instructive cues from their final cortical circuit niche.

Interneurons migrate long distances prior to circuit integration and must devote a significant part of their transcriptional program to maintain the molecular machinery needed for migration (Cobos et al., 2007; Peyre et al., 2015). During this process, which entails leaving local cues behind, their identity must remain intact until they reach their final location, regardless of extrinsic signals they may encounter along their path. This could be achieved through the control of transcriptional programs or through epigenetic mechanisms. The latter has been shown to play a role in the identity of interneurons migrating towards the olfactory bulb (Banerjee et al., 2013). But it is unclear which transcriptional programs, if any, might be necessary to maintain cortical interneuron identity during this process.

All MGE-born interneurons express the transcription factor Sox6, which is continuously expressed during migration and postnatal maturation (Batista-Brito et al., 2009). Complete early postmitotic deletion of Sox6 in the mouse disrupts laminar positioning of Sst^+^ and Pvalb^+^ neurons and maturation of the latter class in particular (Azim et al., 2009; Batista-Brito et al., 2009). Importantly, Sox6 has also been extensively implicated in cell fate determination and differentiation in other cellular systems (Hagiwara, 2011). Its action is dependent on its interaction with additional effector proteins (Kamachi et al., 2000; Lee et al., 2014), providing Sox6 with the molecular versatility to affect a variety of systems at multiple developmental stages (Lefebvre, 2010; Panman et al., 2014; Stolt et al., 2006).

Here we show that migrating Sst^+^ neurons require Sox6 expression specifically during migration and initial network integration in order to maintain their previously acquired subtype identity. Combining scRNAseq, *in situ* hybridization and electrophysiology, we show that loss of Sox6 during migration results in reduced subtype diversity but not in the survival or migration of Sst^+^ cells, indicating that maintenance of Sst^+^ interneuron subtype identity during migration relies on an active transcription factor program.

## Results

### Sox6 removal in migrating Sst^+^ neurons does not lead to cell loss nor migration deficits

While still in the MGE subventricular zone, postmitotic interneurons possess core elements of their mature molecular subtype identity (Mayer et al., 2018; Mi et al., 2018). Therefore, migrating interneurons must maintain this molecular identity throughout their migratory journey, regardless of extrinsic cues they may encounter along the migratory route or within their final cortical niche. The mechanism by which this maintenance occurs is unknown.

Sox6 expression begins around when MGE-derived cells exit the cell cycle in the MGE (Batista-Brito et al., 2009). We utilized Sst^cre^;Rosa-Cag-Egfp;Sox6^fl/±^ mice to specifically remove Sox6 in migrating Sst-expressing interneurons, while labelling affected cells with eGFP (Figure 1A). SstCre expression initiates only after the cells have started their tangential migration (Favuzzi et al., 2019; Taniguchi et al., 2011) a point at which they already express marker genes as exemplified with *Chodl* (Figure 1B). Immunofluorescence analysis at postnatal day (P)21 revealed a nearly complete removal of Sox6 in eGFP^+^ (Sst-expressing) neurons (Figure 1C, D; Control: 92.8 ± 1.6%; Sox6-cKO 4.4 ± 1.1% of eGFP^+^Sox6^+^; p < 0.001; n = 4 per genotype). Importantly, Sox6 loss during migration did not affect the number of cells reaching the primary somatosensory cortex (S1) or their laminar distribution, as assessed by number of eGFP-positive cells per cortical layer (Figure 1C, quantification in 1E). This is strikingly different from the effects of early removal using the Lhx6Cre driver line, which leads to ectopic distribution of MGE-derived cells (Batista-Brito et al., 2009). This suggests that after Sst^+^ interneurons start migrating, layer allocation is no longer regulated by Sox6 expression.

**Figure 1.**
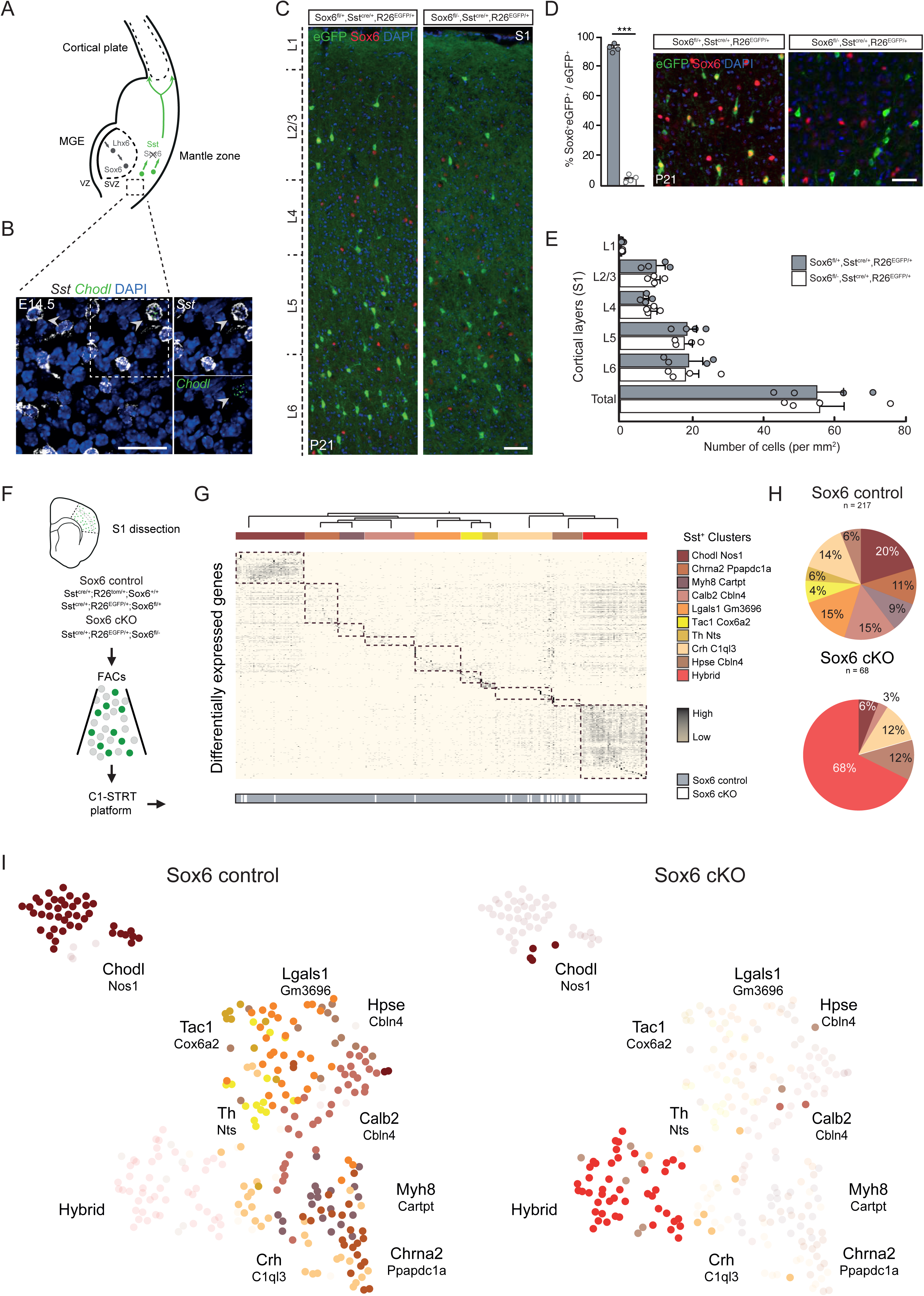
ScRNAseq reveals loss of Sst subtype identity after Sox6 removal during migration. (A) Sst promoter-driven Cre recombinase expression in interneurons initiating tangential migration. (B) *In situ* hybridization in E14.5 brains shows neurons co-expressing *Sst*^+^*Chold*^+^ after exiting the MGE (Scale bar: 25 μm). (C) Immunofluorescence for eGFP and Sox6 in S1 coronal sections of P21 *Sst*^*cre/*+^;*Rosa-Cag-Egfp;Sox6*^*fl±*^ mice (Scale bar: 50 μm). (D) Quantification of eGFP^+^ cells expressing Sox6 in control and Sox6-cKO at P21 (n = 4 per genotype) Scale bar: 25 μm. (E) Laminar distribution of eGFP^+^ per mm^2^ (n = 4 per genotype). (F) Schematic representation of experimental workflow (minimum of 4 animals were used per group). (G) Heatmap displaying 500 mostly differentially expressed genes in clustered cells after BACKspin analysis. Each column represents a cell and each row, a gene. Bottom: in grey, 217 control cells; in white, 68 Sox6-cKO cells. (H) Pie charts display percentages of each subtype in the two groups. (I) Left t-SNE highlights control cells; right t-SNE highlights Sox6-cKO cells. t-SNEs using dataset in F. MGE: medial ganglionic eminence; S1: primary somatosensory cortex; SVZ subventricular zone; VZ: ventricular zone. p-value < 0.001(***). Error bars = standard error of the mean (SEM).

### Sox6 is required for maintenance of subtype identity in Sst^+^ neurons

Sox6 has been linked to cell type specification (Hagiwara et al., 2005; Hagiwara et al., 2007) and differentiation in several systems (Stolt et al., 2006), as well as neuronal cell development (Panman et al., 2014). Through interaction with various co-factors, Sox6 can regulate unique gene networks in different cellular contexts and neuronal subtypes (Hagiwara, 2011; Taglietti et al., 2016). Therefore, in light of the extensive heterogeneity of the Sst^+^ interneuron class, we sought to investigate whether Sox6 played a role in maintaining the identity of these neurons.

To determine this putative role, we sorted tdTomato-or GFP-expressing cells from dissociated S1 cortical slices collected from control and conditional Sox6 mice between P21 and P28 to perform scRNAseq (Figure 1F). After quality control and identification of significantly differentially expressed genes (see methods), we clustered 217 control and 68 Sox6-cKO cells based on their molecular identity. Our scRNAseq profiling was utilized as a platform to assess the cells’ entire transcriptome and subsequently validate our findings in tissue.

This analysis revealed a total of ten distinct molecular subtypes (Figure 1G). In a recent study, 15-30 Sst-expressing subtypes were described, depending on cluster stringency (Tasic et al., 2018). Control cells were distributed amongst nine distinct clusters (Figure 1G,H) and each cluster expressed key molecular markers corresponding to a specific counterpart or to a fusion between closely related clusters previously described (Tasic et al., 2018): Chodl-Nos1 (corresponding to *Sst-Chodl*), Chrna2-Ppapdc1a (*Sst-Chrna2* clusters), Myh8-Cartpt (*Sst-Myh8* clusters), Calb2-Cbln4 (*Sst-Calb2-Pdlim5*), Lgals1-Gm3696 (*Sst-Crh2r, Sst-Esm1*), Tac1-Cox6a2 (*Sst-Tac1* clusters), Th-Nts (*Sst-Nts*), Crh-C1ql3 (*Sst-Tac2-Tacstd2, Sst-Rxfp1*), Hpse-Cbln4 (*Sst-Hpse* clusters). Strikingly, five of the nine Sst-expressing subtypes found in the control were entirely absent from Sox6-cKO cells and two others were drastically reduced (Figure 1G-I). Specifically, the following subtypes were absent: Chrna2-Ppapdc1a, Myh8-Cartpt, Lgals1-Gm3696, Tac1-Cox6a2, Th-Nts, while only few cells were identified as Chold-Nos1 and Calb2-Cbln4 (Figure 1G,H). Remarkably, the Sst-expressing subtypes in the Sox6-cKO cortex comprised two molecular subtypes found in the control, Crh-C1ql3 and Hpse-Cbln4, and a third exclusive cluster that did not include any control cells, which seemed to be a molecular hybrid of these subtypes (Figure 1G-I).

To confirm our scRNAseq findings, we selected subtype specific markers to identify specific Sst cell types (Figure 2A). We performed *in situ* hybridization targeting four (of seven) subtypes absent or reduced in Sox6-cKO S1 cortex (Sst^cre^;Sox6^fl/-^) (Figure 2B-E). Chodl-Nos1 neurons are well-characterized long-range projecting GABAergic neurons that comprise a consistently unambiguous molecular cluster in scRNAseq studies (Tasic et al., 2016; Tasic et al., 2018; Tomioka et al., 2005; Zeisel et al., 2015). As previously described, the majority of control *Sst*^+^*Chodl*^+^ neurons were found in layer (L)2/3 and L6 (Perrenoud et al., 2012; Tomioka et al., 2005), while in the Sox6-cKO cortex these cells were virtually absent in all layers (Figure 2B, Control, 4.3 ± 0.3%; Sox6-cKO, 0.7 ± 0.4% of Sst^+^ cells, p = 0.002, n = 3 per genotype). Two other subtypes investigated were exclusively located in deep layers and were also reduced in the Sox6-cKO cortex (Figure 2C,D): Sst^+^Chrna2^+^ (Control, 7.4 ± 1.7%; Sox6-cKO, 1.4 ± 0.7% of Sst^+^ cells, p = 0.03, n = 3 per genotype) and Sst^+^Th^+^ (Control, 4.1 ± 0.9%; Sox6-cKO, 1.4 ± 0.4% of Sst^+^ cells, p = 0.058, n= 3 per genotype). Sst^+^Tac1^+^ cells (Figure 2E) corresponded to a small percentage of all Sst^+^ cells (Control, 2.7 ± 0.4%; n = 3). In Sox6-cKO, only Sst^+^Tac1^+^ cells in L5 were significantly reduced (L5: Control, 1.4 ± 0.4%; Sox6-cKO, 0.0 ± 0.% of Sst^+^ cells, p = 0.02, n = 3 per genotype).

**Figure 2.**
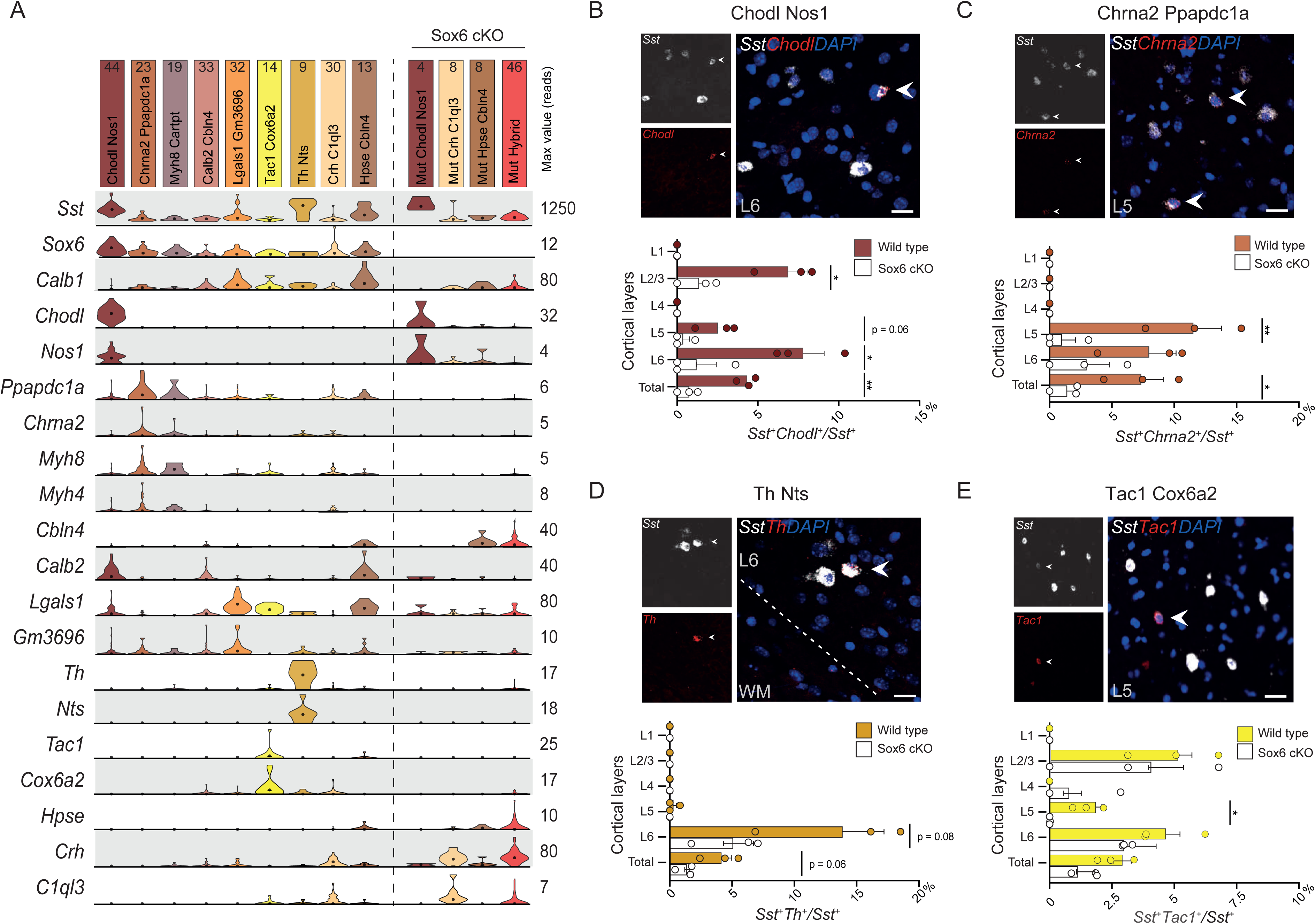
Loss of subtypes confirmed by *in situ* hybridization. (A) Violin plots show the expression of key subtype markers, dots represent the median value. (B-E) Representative *in situ* hybridization of key markers for four distinct subtypes in wild type S1, co-expressed with *Sst*. Bar graphs show the layer distribution of the distinct subtypes in wild type *versus* Sox6-cKO cortex (n=3 per group). p-value < 0.05 (*), < 0.01 (**), < 0.001 (***).Scale bar 25 μm. Error bars = SEM.

### Molecular and electrophysiological characterization of Sox6-cKO Sst^+^ neurons

Loss of Sox6 does not affect the number of Sst^+^ cells nor layer location but leads to loss of Sst^+^ subtype diversity. Describing Sst^+^ neurons from a perspective of hierarchical identity, we compared Sox6-cKO cells to control cells with regards to the expression of genes corresponding to distinct segments of neuronal identity (Figure 3A). Sox6-cKO cells expressed pan-neuronal markers, GABA-producing machinery, as well as Sst-lineage specific markers (including *Sst, Satb1, Elfn1, Lhx6, and Mafb*; Figure 3A). Nevertheless, genes related to their refined subtype identity were affected upon Sox6 removal. We observed a loss of specificity in the expression of marker genes including *Hpse, Crh, C1ql3, Cbln4, Calb2,* and *Pvalb,* particularly within the hybrid cluster (Figure 3A).

**Figure 3.**
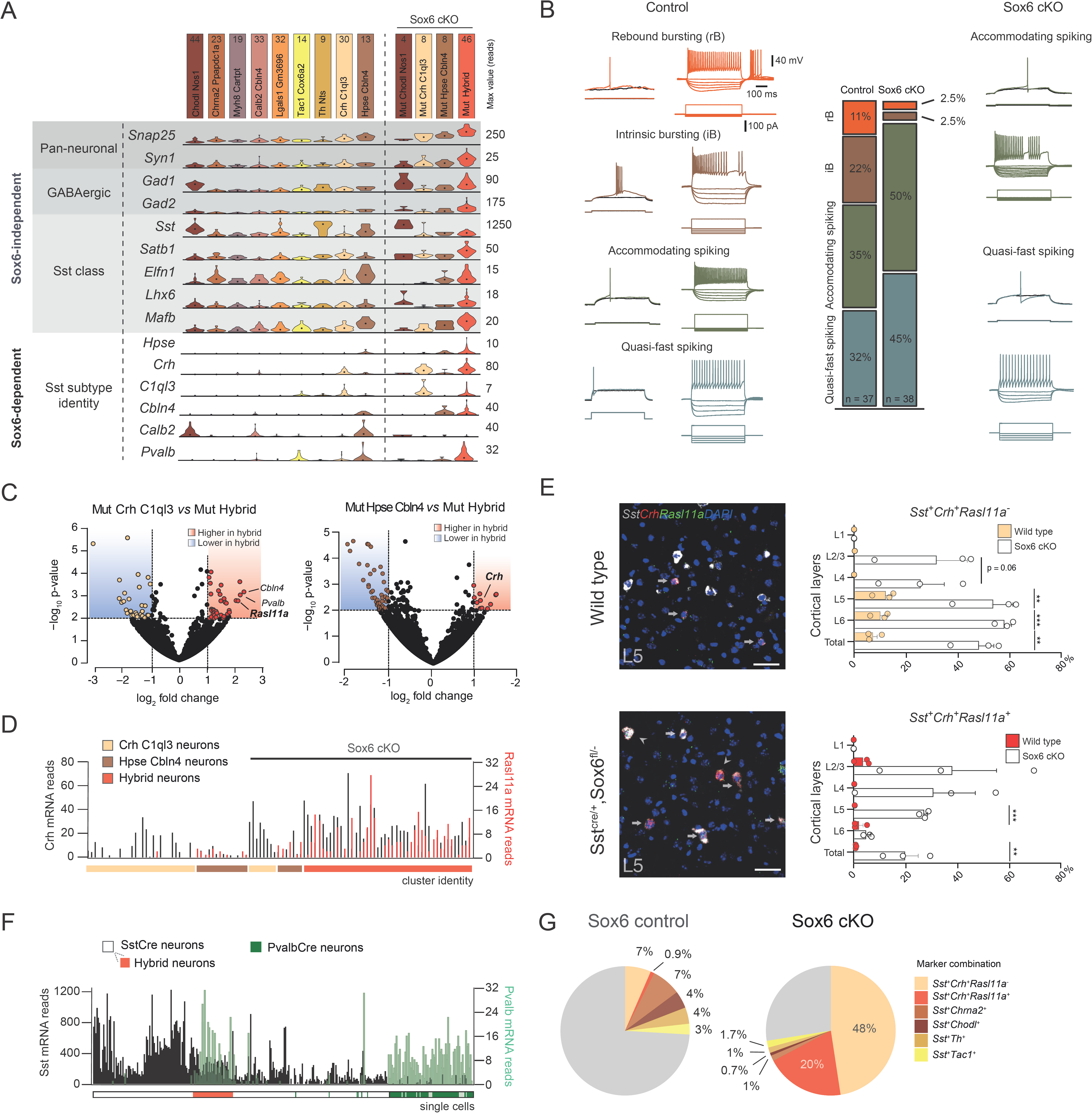
Sox6-cKO cells retain characteristic molecular and electrophysiological properties of Sst-neuron class. (A) Violin plots show mRNA expression of key markers associated with distinct identity-acquiring steps; dots represent the median value. (B) Whole-cell patch-clamp recording of L5 Sst-expressing neurons in *Sst*^*cre/*+^;*Rosa-Cag-Egfp;Sox6*^*fl±*^ reveals that Sox6-cKO cells have intrinsic properties comparable to control cells. Rebound and intrinsic bursting neurons are nearly absent from the Sox6-cKO cortex. (C) Volcano plots (DESeq2) show differentially expressed genes comparing hybrid cells *versus* Mut Crh-C1ql3 or Mut Hpse-Cbln4. X-axis dashed line: p-value < 0.01. Y-axis dashed line: log2 fold change −1 < and >1. (D) Number of *Crh* (black) and *Rasl11a* (red) mRNA reads in Crh-C1ql3, Hpse-Cbln4, and hybrid neurons. (E) Representative *in situ* hybridization of S1 L5 for Sst, *Crh*, and *Rasl11a*, in both wild type and Sox6-cKO. Full arrows indicate Sst^+^Crh^+^Rasl11a^-^ cells (Crh-Cq1l3 cells), while arrowheads indicate Sst^+^Crh^+^Rasl11a^+^ (hybrid cells) cells, found only in Sox6-cKO cortices. Bar graphs show their layer distribution in wild type *versus* Sox6-cKO cortex (n=3 per genotype). (F) Pie charts display percentages of the Sst^+^ cell types investigated (distinct marker combinations). (G) *Sst* (black) and *Pvalb* (green) mRNA expression in sorted SstCre and PvalbCre neurons (bottom: each line represents a single cell, SstCre in white, including hybrid cells in red, and PvalbCre cells in green). p-value < 0.01 (**), < 0.001 (***). Scale bar 50 μm. Error bars = SEM.

Accordingly, whole-cell patch clamp recordings in L5 neurons in S1 revealed that Sox6-cKO neurons exhibited electrophysiological properties within the normal range of wild-type Sst-neurons (Figure 3B, Supplementary Table 1). However, while control cells showed diverse electrophysiological profiles including accommodating spiking cells, rebound bursting (exhibiting a burst of action potentials after a hyperpolarizing current), intrinsic bursting (bursting from action potential threshold) neurons, as well as and the characteristic quasi fast-spiking pattern previously described (Nigro et al., 2018; Xu et al., 2013), Sox6-cKO cells displayed less diversity. Recordings from Sox6-cKO neurons revealed that rebound bursting neurons were lost (control 11%, 4/37 *versus* Sox6-cKO 2.5% 1/38; Figure 3B). Rebound bursting properties have recently been shown to be characteristic of *Chrna2*-expressing Sst neurons, a specific T-type Martinotti cell type enriched in deep layers (Hilscher et al., 2017). Loss of rebound bursting in Sox6-cKO cells thus tightly corresponds to our transcriptomic results in which *Chrna2*-expressing cells are absent in the Sox6-cKO cortex (Figure 2C). In addition to the lack of rebound bursting cells, we did not observe intrinsically bursting cells previously described (Butt et al., 2005). At last, quasi-fast spiking cells constituted 45% of the L5 Sox6-cKO cells recorded (Figure 3B). Also known as X94 cells, these neurons have been recently shown to be labeled by *Hpse* (Naka et al., 2019), which indeed corresponds to one of the remaining Sst^+^ cell types revealed by scRNAseq (Figure 1G). Comparing intrinsic and firing properties of control accommodating spiking cells and X94 cells with their Sox6-cKO counterparts revealed normal electrophysiological properties in the Sox6-cKO cells (Supplementary Table 1), despite decreased diversity of electrophysiological profiles due to the loss of other subtypes.

A majority of the Sox6-cKO cells (68%) analyzed with scRNAseq formed a distinct cluster of hybrid neurons that shared markers belonging to the two remaining subtypes: Crh-C1ql3 and Hpse-Cbln4 (Figure 1H and 2A). In order to confirm our findings from scRNAseq and reveal the location of the hybrid cell subtype, we selected markers to distinguish hybrid cells from the remaining subtypes. Pairwise differential expression analysis (using DESeq2; (Love et al., 2014) between hybrid cells *versus* Sox6-cKO Crh-C1ql3 or Sox6-cKO Hpse-Cbln4 revealed that the co-expression of *Sst, Crh* and *Rasl11a* distinguishes hybrid cells from the other two subtypes (Figure 3C). Sox6-cKO Crh-C1ql3 cells expressed *Sst* and *Crh* but not *Rasl11a* while hybrid cells express all three markers (Figure 3C). Closer inspection of the scRNAseq data revealed that we did not detect *Crh* transcripts in all control Crh-C1ql3 cells (Figure 3D), though all Sox6-cKO Crh-C1ql3 cells expressed it. If this was a true signal and not due to false drop-outs, this could lead to an underestimation of the number of Crh-C1ql3 cells in the control brains.

Nonetheless, *in situ* hybridization revealed that *Sst*^+^*Crh*^+^*Rasl11a*^*-*^ cells were located exclusively in L5-6 in control animals, whereas in the Sox6-cKO this analysis revealed a drastic increase in the number of *Sst*^+^*Crh*^+^*Rasl11a*^*-*^ cells (Control: 6.5 ± 0,2%; Sox6-cKO: 48 ± 5.6% of Sst^+^ cells, p = 0.002, n = 3 per genotype) which occupied L2-6 (Figure 3E). Furthermore, we detected very few cells co-expressing *Rasl11a, Sst* and *Crh* in the control cortex, while in the Sox6-cKO cortex of 20% of all Sst-expressing cells co-expressed the three markers (Control: 0.92 ± 0.2%; Sox6-cKO: 20 ± 5.1% of Sst^+^ cells, p = 0.02, n = 3 per genotype) (Figure 3E). Therefore, while only 8% of Sst-expressing neurons express *Crh* in controls, our *in situ* analysis demonstrates that 68% of all Sst-expressing neurons express *Crh* in Sox6-cKO S1 (Figure 3F).

Finally, our scRNAseq data revealed that the Sox6-cKO hybrid cluster expresses high levels of *Pvalb* mRNA (Figure 3A,C). Because Sox6 plays a crucial role in Pvalb-expressing neuron maturation (Batista-Brito et al., 2009), we sorted wild type PvalbCre cells and processed them for scRNAseq to address whether Sox6-cKO cells’ identity pivoted towards a Pvalb-expressing one. Clustering of PvalbCre cells with all control and Sox6-cKO Sst-expressing cells revealed that PvalbCre cells clearly clustered separately from Sox6-cKO hybrid Sst^+^ neurons, arguing that loss of Sox6 in Sst-expressing neurons does not lead to a conversion into classical Pvalb-expressing interneurons (Figure 3G).

### Time-dependent action of Sox6 in Sst^+^ neurons

Our results show that Sox6 is necessary to maintain the subtype diversity of Sst^+^ neurons during migration. While virtually all Sst^+^ neurons expressed Sox6 at P21 (Figure 1 C,D), at P90 only 33 ± 8% of all eGFP^+^ (Sst-expressing) neurons express Sox6 (60% reduction, p < 0.001, n = 3 per genotype) (Figure 4A). This suggests that Sox6 might not be necessary for the maintenance of subtype identity once Sst^+^ neurons are mature. One of the subtypes severely affected by Sox6 removal during migration was the Chodl-Nos1 subtype (Figure 2B). To assess if Sox6 is necessary to maintain the identity of Chodl-Nos1 cells after migration has been completed, we conditionally removed Sox6 at the end of the first postnatal week, a time when cells have completed migration and begun forming synaptic connections (Chattopadhyaya et al., 2004; Favuzzi et al., 2019). For that, we utilized the tamoxifen (tx) inducible pan-interneuronal transgenic mouse line Dlx1/2CreERT (Figure 4B). Tx injections at P7 followed by immunofluorescence analysis at brains collected at P14 (TxP7-14) show that Sox6 is efficiently removed from eGFP^+^ cells (Control, 89 ± 2%; Sox6-cKO, 6 ± 0.8%, p < 0.001, n = 3 per genotype). In order to identify the Sst-Chodl-Nos1 population only amongst eGFP-expressing cells, we performed immunofluorescence for eGFP, Sst and Nos1, a marker combination unique to this population (Figure 2A). When brains were analyzed at P28 (TxP7-P28, Figure 4C), the percentage of cortical eGFP^+^Sst^+^Nos1^+^ cells was intact (Control, 3.1 ± 0.2%; Sox6-cKO, 2.9 ± 0.1% of all eGFP^+^ cells, p = 0.53, n = 4 and 3 respectively). This suggests that after circuit integration Sst^+^ neurons no longer rely on the same Sox6-dependent mechanism to maintain their subtype identity.

**Figure 4.**
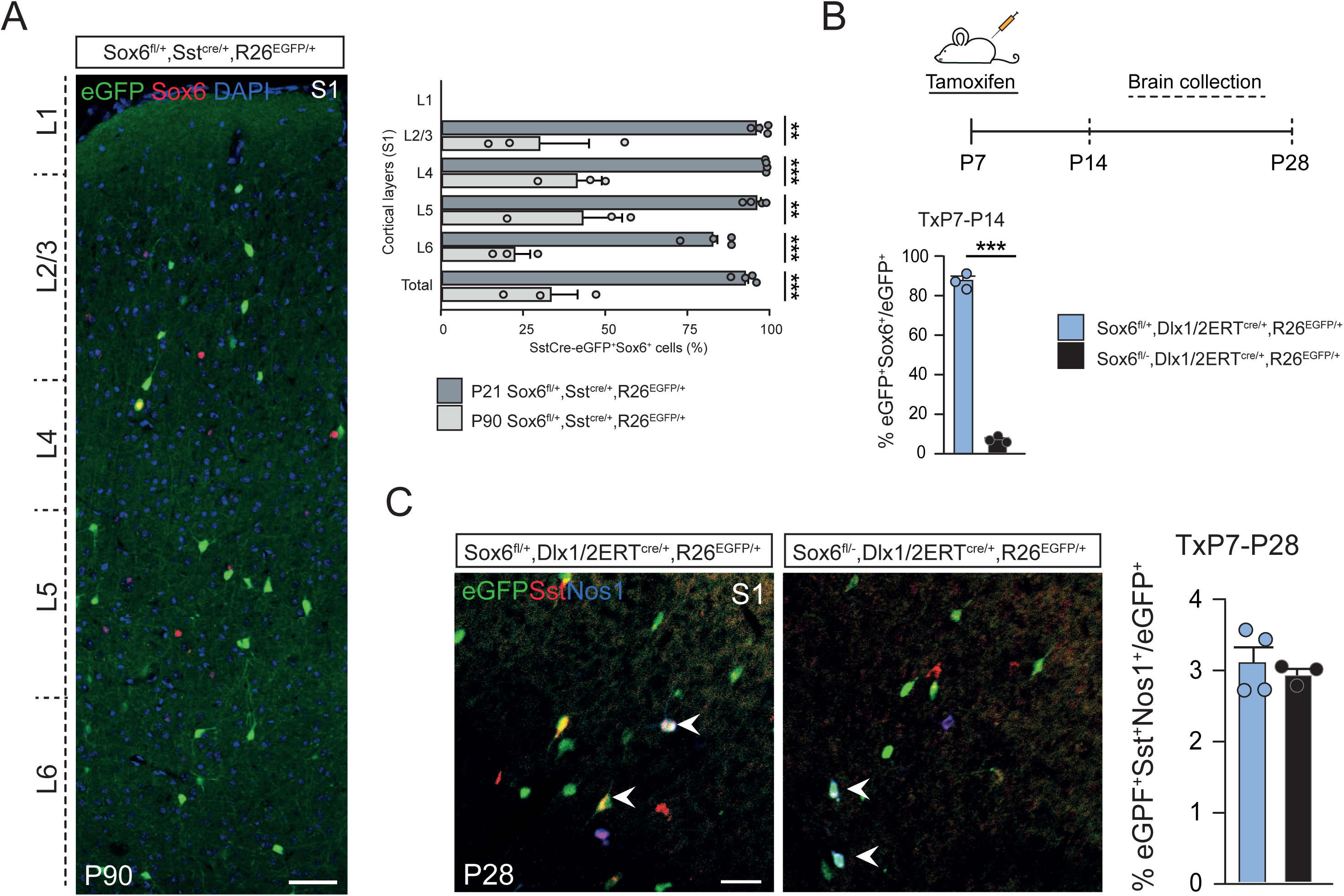
Postnatal removal of Sox6 does not disrupt Chodl (Sst^+^Nos1^+^) expression. (A) Left: S1 coronal section of P90 Sst^cre/+^;Rosa-Cag-Egfp;Sox6^fl/+^ mouse. Right: Quantification of eGFP^+^ cells expressing Sox6 in P21 and P90 control animals (n = 4 and 3 respectively). (B) Top: experimental timeline representing tamoxifen (tx) injections at P7 and brain collection at P14 or P28. Bottom: quantification of eGFP^+^ cells expressing Sox6 in control and Sox6-cKO at P14 (n = 3 per genotype). (C) Quantification of immunohistochemistry of eGFP^+^ cells co-labelled with Sst and Nos1 (markers combination unique to Chold-Nos1 cells). Tx: tamoxifen. p-value < 0.001 (***). Scale bar 50 μm. Error bars = SEM.

## Discussion

Here we show that loss of Sox6 in Sst-expressing interneurons during tangential migration altered Sst-subtype identity without affecting the expression of Sst-class identity genes or the development of mature electrophysiological features. These findings suggest that the period between initial specification and network integration is a conceptually (and molecularly) discrete time window in the life of an interneuron.

Mitotic and/or early postmitotic expression of Sox6 is known to be necessary for the induction of Sst^+^ cells and correct migration of interneurons (Azim et al., 2009; Batista-Brito et al., 2009), and here we show that Sox6 has an additional role in subtype maintenance during migration but not after network integration.

Fate restriction in MGE interneuron progenitors begins in response to sonic hedgehog, which induces and maintains *Nkx2-1* (Gulacsi and Anderson, 2006; Xu et al., 2005). *Nkx2-1* is necessary to establish and maintain early MGE progenitors (Butt et al., 2008). While still in the MGE, *Nkx2-1* induces *Lhx6* expression which is necessary for the acquisition of general MGE interneuron identity and later maturation of Sst-and Pvalb-expressing cell types (Liodis et al., 2007; Zhao et al., 2008). Sox6 has been implicated in the levels of nNOS and Sst expression in Sst-interneurons (Jaglin et al., 2012) but our data offer an alternative explanation in that Sst-Chodl-Nos1 neurons are losing their subtype but not Sst-identity. Beyond the broad specification of class identity, recent studies suggest that MGE-born interneurons attain the molecular signature of their nuanced adult subtype identity before tangential migration starts (Mayer et al., 2018; Mi et al., 2018) (Figure 1B). Even if the cellular diversity observed within the MGE does not parallel the full molecular heterogeneity of the adult cortex (Codeluppi et al., 2018; Tasic et al., 2018; Zeisel et al., 2018), several Sst-expressing subtypes can be identified within embryonic postmitotic neurons (Mayer et al., 2018; Mi et al., 2018), including the subtypes expressing Chodl (named Chodl-Nos1 in our study) and Myh8 (named here, Chrna2-Ppapdc1a), which were specifically lost in the Sox6-cKO.

During tangential migration different Sst^+^ subtypes can also be identified after they have entered different migratory routes (e.g. Sst^+^Cbln4^+^ in the mantle zone and Sst^+^Chodl^+^, in the subventricular zone) (Lim et al., 2018). Altering their migratory path by deleting the transcription factor *Mafb* only affects minor morphological aspects of their final identity, leaving electrophysiological and key molecular markers intact (Lim et al., 2018). This is consistent with an identity commitment prior to arriving in the developing cortex and less so with a model where cells read out identity-inducing signals as they migrate towards their final destination. The former model is further supported by heterotopic, isochronic transplantations of interneurons between cortex and hippocampus suggesting that the Sst-Chodl (nNos-expressing) cells keep their donor-identity (Quattrocolo et al., 2017). We therefore consider it unlikely that the effect that we see on certain subtypes of Sst^+^ cells reflects different requirements with regards to extracellular cues during migration. While our data do not exclude the possibility that Sox6 could play a role in the nuanced specification of the other Sst^+^ subtypes, this strongly suggests that Sox6 was necessary to maintain the Chodl-Nos1 and Chrna2-Ppapdc1a identity during migration rather than being necessary for their finer specification.

An important element of our findings is that, by maintaining distinct Sst^+^ subtype identities, Sox6 is acting in distinct cellular contexts to sustain unique transcriptomic profiles. This poses the question of how the same shared transcription factor can differentially activate and/or repress gene sets in specific subtypes of Sst^+^ neurons. In the nematode *C. elegans*, unc-3 is a transcription factor with a similar role: all six subtypes of ventral cholinergic motor (VCM) neurons express unc-3, which is able to initiate and maintain the distinct neuronal features of each neuronal type, by co-acting with complementary, class-specific repressing transcription factors (Kerk et al., 2017; Kratsios et al., 2011). Similar to the effect of Sox6 ablation in cortical Sst^+^ neurons, loss of unc-3 disrupts VCM neuronal class diversity and generates a “hybrid” motor neuron (Kerk et al., 2017; Kratsios et al., 2011). Our findings in mammalian cortical interneurons suggest this to be a conserved developmental strategy to generate cellular diversity. Consistent with this idea, Sox6 is capable of to both activating and repressing specific gene sets through interaction with different partner effectors (Hagiwara et al., 2005; Hagiwara et al., 2007; Liu et al., 2007; Taglietti et al., 2016). It acts primarily by binding to enhancers and super-enhancers (Kawane et al., 2014; Liu and Lefebvre, 2015), which are regions of DNA with abundant transcription factor binding whose activation is particularly associated with genes important for cell identity (Achour et al., 2015). In the context of Sst-expressing neurons, Sox6 appears to act in two distinct manners. First, since Sox6-cKO hybrid cells shared markers common to Crh-C1ql3 and Hpse-Cbln4 cell types, Sox6 may act to repress genes important for the final differentiation of these subtypes. Secondly, since Sox6-cKO cells lacked markers associated with any of the absent subtypes (*Chrna2, Chodl, Tac1* etc.), Sox6 is likely to also act as co-activator of such molecular programs. In other words, our results corroborate a dual repressor/activator role for Sox6 in Sst^+^ interneurons.

Numerous transcription factors involved in the early steps of cell identity formation continue to be expressed in postmitotic neurons to maintain that identity (Deneris and Hobert, 2014). Interestingly, a complete knock-out of Sox6 leads to a severe loss of Sst expression (70% decrease) and disturbed layer distribution in the absence of GABAergic cell loss (Azim et al., 2009). Therefore, it is possible that early Sox6 expression plays a role in determining the subtype identity of mitotic MGE-born cells. Conversely, loss of Sox6 just as the MGE-cells exit the cell cycle (using Lhx6Cre) resulted in only a 15% decrease of Sst expression and perturbed layer distribution. Our data suggest that Sox6 has yet another later distinct role in subtype-identity maintenance during migration, particularly for Chodl-Nos1 cells.

In the mouse, the term “master regulator” has been used to describe transcription factors necessary for the initiation of entire class-identity programs - e.g. Nkx-2.1 in MGE-derived interneurons (Butt et al., 2008). Nkx-2.1 is, however, downregulated prior to the cells entering the dorsal pallium (Nobrega-Pereira et al., 2008) and is therefore not necessary for maintenance of these cells’ identity throughout life. Early gene-removal experiments (mitotic or just after cell cycle-exit) have implicated other genes in the induction of Sst^+^ interneuron identity and maturation, including *Lhx6, Satb1, Sp9, Mafb* and *Dlx*-genes (Close et al., 2012; Cobos et al., 2007; Denaxa et al., 2012; Du et al., 2008; Liodis et al., 2007; Liu et al., 2018; Pai et al., 2019; Vogt et al., 2014; Zhao et al., 2008). Additionally, late removal of *Dlx1/2* affects interneuron migration, network integration and other functional aspects such as GABA-levels (Pla et al., 2018). However, none of these genes have been implicated in the maintenance of identity. While in nematodes, the term “terminal selector gene” has been used to describe genes that not only induce but also maintain cell identity, to our knowledge no gene that fits the formal definition of a terminal selector gene has been experimentally shown in mammals. In mouse, *Lhx6* (which is upstream from *Sox6*; (Batista-Brito et al., 2009) plays a role in both cell-specification and the survival of neuronal cell types, as shown by manipulation of gene-dosage through heterozygous removal (Neves et al., 2013). However, in order to argue for a definite role in adult neurons, one needs to rule out subtle effects on cell-specification, that can explain the later phenotypes. Thus, the gene in question must be removed after normal generation and cell type-specification has occurred to specifically test a late function. Our data suggest that mammalian neurons utilize a bridge-program (*i.e.* that is then later lost), such as that described here for Sox6, to maintain initial subtype identity specifically through the migratory period and into circuit integration.

## Acknowledgements

The authors would like to thank the Eukaryotic Single Cell Facility at SciLifeLabs for support, Véronique Lefebvre for providing the Sox6 CKO mouse line, Gordon Fishell for Dlx1/2CreERT, Sst-cre and RCE mouse lines and Michael Wegner for providing Sox6 antibody. H.M. was funded by the KI doctoral program. J.H.-L. was funded by the Swedish Research Council (Vetenskapsrådet, awards 2010-3103 and 2014-3863), StratNeuro, Jeanssons Stiftelser and the Swedish Brain Foundation (Hjärnfonden).

## Author contributions

Conceptualization, H.M. and J.H-L., Methodology; H.M., H.H., R.B-B., A.B.M-M., S.L., J.H-L; Investigation: H.M., K.N., H.H., P.O., A.K., J.R., and J.H-L.; Formal Analysis: H.M., K.N., H.H., Resources: S.L., J.H-L.; Writing Original Draft; H.M., J.C., and J.H-L; Writing – Review and Editing; All authors. Visualization: H.M.; Supervision: J.H-L.

## Methods

### Experimental animals

All animal handlings were according to local ethical regulations and were approved by the local committees for ethical experiments on laboratory animals (Stockholms Norra Djurförsöksetiska nämnd, Sweden). We utilized Sst-IRES-Cre (The Jackson Laboratory, Stock No: 013044; (Taniguchi et al., 2011) crossed with R26R CAG-boosted EGFP (RCE) (The Jackson Laboratory, Stock No: 32037; (Miyoshi et al., 2010)or a Rosa26-tdTomato strain, together with a combination of Sox6 loxp/deletion background (Mouse Genomic Informatics, MGI:3641204, MGI:3641205; (Dumitriu et al., 2006) combined to generate Sst^cre/+^;Rosa-Cag-Egfp;Sox6^fl/±^ animals. We also used the tamoxifen (tx) inducible pan-interneuronal Dlx1/2-Cre-ERT2 (The Jackson Laboratory, Stock No: 014600) crossed with the same reporter and Sox6 loxp background. Tx was diluted in corn oil (20 mg/ml) and approximately 0.2 ml were administered per pup (P7) and brains were collected at P14 or P28 (n= 3 to 5 per condition), following procedures on sections bellow

### Single cell preparation

P21-28 brains were collected and processed following the same protocol as that used for acute-slice electrophysiology (for details, see section Slice Electrophysiology). Tissue corresponding to S1 was dissected and dissociated into a single cell suspension, as previously described (Munoz-Manchado et al., 2018). The tissue was then dissociated using the Papain dissociation system (Worthington) following the manufacturer’s instructions. Cell suspension obtained was filtered with 20 μm filter (Partec) and kept in cold oxygenated HBSS solution (Sigma) with 0.2% BSA and 0.3% glucose. Cells were sorted based on fluorescence (eGFP or tdTomato) prior to running them on the C1-STRT platform, followed by a subsequent manual loading into the C1 chip (Fluidigm system) and standard sample processing: lysis, reverse transcription, PCR, tagmentation, high-throughput sequencing (Illumina) and molecule counts (Munoz-Manchado et al., 2018). Doublets were excluded and only fluorescent single cells were processed for RNA sequencing.

### Immunohistochemistry

Mice were perfused with ice-cold phosphate buffered saline (PBS) followed by 4% paraformaldehyde (PFA) in PBS. Dissected brains were further fixed for 1 hour in 4% PFA at 4°C and cryoprotected by incubation in 30% sucrose/PBS at 4°C overnight. Brains were then embedded in optimal cutting temperature (OCT, Histolab Products AB) and stored at −80°C until further processing. Ten to 14 µm sections were cut using a cryostat (Cryostar NX70), washed with PBST (PBS containing 0.1% Tween20), and incubated in a blocking buffer (PBS containing 2.5% normal goat serum, 2.5% donkey serum, 2.5% BSA, 0.5 M NaCl, and 0.3% Tween 20) for 1 hour at room temperature (RT). Sections were then incubated with primary antibodies in dilution buffer (2.5% BSA, 0.5 M NaCl, and 0.3% Tween 20 in PBS) overnight at 4°C, followed by 3 washes in PBS for 15 minutes each, and incubation with secondary antibodies for 1 hour at RT. Sections were rinsed 4 times in PBS for 10 min each and nuclei were stained by incubation with 100 ng/mL 4,6-diamidino-2-phenylindole (DAPI) in PBS for 5 minutes at RT.

The following primary antibodies were used: chicken anti-green fluorescent protein (1:2000, Abcam), guinea pig anti-Sox6 (1:2000, not commercially available), rat anti-Sst (1:500; Millipore), goat anti-Nos1 (1:2000; not commercially available). For visualization of signal, secondary antibodies conjugated with Alexa Fluor dyes 488 (1:1000; Invitrogen), 555 (1:400; Invitrogen) and 647 (1:400; Invitrogen) were used. Sections were then coverslipped using Fluoromount-G (Southernbiotech).

### *In situ* hybridization

Brains from P21-30 mice, three wild-type CD1 and three Sox6-cKO (SstCre:Sox6^fl/-^), were collected and freshly frozen in OCT compound (Tissue-tek) on dry ice and kept at −80°C until sectioning. Coronal sections with 10 µm thickness were obtained with a cryostat (Cryostar NX70),and kept in −80°C until use. *In situ* hybridization was performed with RNAscope technology (Advanced Cell Diagnostics Biotechne) according to the manufacturer’s instruction, with probes targeting following genes: *Sst, Chodl, Th, Tac1, Chrna2, Rasl11a, Crh*. For E14.5 samples, brains were fixed with PFA 4% for 4 hr prior to sectioning (10 µm). *In situ* hybridization was performed following the manufacturer’s instructions.

### Imaging and cell quantification

Tile scan images spanning all the six layers of S1 were acquired on a Zeiss LSM 700 or LSM 800 confocal microscope equipped with a Plan Apochromat 10x, 20x and 40x objectives. Cortical layers were determined by the DAPI signal density and cells were counted using the Cell Counter plugin in ImageJ (U.S. National Institutes of Health).. All data are presented as mean ± standard error of the mean (SEM

### Slice electrophysiology

Whole-cell patch-clamp electrophysiological recordings were obtained from eGFP-expressing cells in acute brain slices prepared from P12 to P30 Sst^cre/+^;Rosa-Cag-Egfp;Sox6^fl/±^ animals. Animals were anesthetized with ketamine/xylazine (4:1), the brain was quickly removed and transferred to ice-cold cutting-solution of the following composition (in mM): 87 NaCl, 75 sucrose, 2.5 KCl, 25 NaHCO_3_, 1.25 NaH_2_PO_4_, 7 MgCl_2_, 1 CaCl_2_, and 10 glucose. Animals older than P21 were transcardically perfused with cutting-solution. The brain was then fixed to a stage and 300 μm slices were cut on a vibratome (VT1200 S, Leica). Slices were then individually transferred into an incubation chamber containing oxygenated artificial cerebro-spinal fluid (aCSF) at 35°C for 30 min followed by at least 30 min at room temperature before recordings. During recording, slices were continually perfused with oxygenated aCSF composed of: (in mM) 125 NaCl, 2.5 KCl, 25 NaHCO_3_, 1.25 NaH_2_PO_4_, 2 MgCl_2_, 2 CaCl_2_, and 10 glucose. Patch electrodes were made from borosilicate glass (resistance 4–8 MΩ; Hilgenberg, GmbH) and filled with intracellular solution containing (in mM): 135 KCl, 10 Na-Phosphocreatine, 10 HEPES, 4 Mg-ATP, 0.3 Na-GTP.

### Clustering analysis

We sorted a total of 335 eGFP^+^ or tdTomato^+^ individual cells from SstCre S1 dissociated cortex. Our exclusion criteria for low-quality cells was total number counts inferior to 5000 reads, excluding mitochondrial, sex (Xist, Tsix, Eif2s3y, Ddx3y, Uty, Kdm5d), and immediate early genes (Fos, Jun, Junb, Egr1). Twenty-nine SstCre cells were excluded at this step, resulting in 304 cells.

The remaining 304 cells went through a 1^st^ round of clustering (BACKspin algorithm; (Zeisel et al., 2015)using 500 most differentially expressed genes. This revealed cells out of the scope of the study, which were excluded for subsequent analysis: one cell was identified as oligodendrocyte and two groups of cells comprised non MGE-derived neurons, corresponding to VIP cells (expressing Vip, Cck, Htr3a, n = 9) and neurogliaform cells (Lamp5, Car4, Npy, Reelin, n = 9). Next, we reclustered 285 cells (217 controls and 68 Sox6-cKO cells) using the 500 most informative genes. After annotating the clusters in light of the current literature (Tasic et al., 2016; Tasic et al., 2018), we ended up with 10 distinct clusters. When performing DESeq2 differential expression analysis, we selected only genes expressed by at least 2 cells, resulting 10942 genes. We utilized *t*-distributed stochastic neighbor embedding (t-SNE) to visualize the complexity of the ten clusters in two dimensions. We used the log2+1 values of the same 500 most differentially expressed genes used when clustering. Wild type PvCre sorted cells (n = 75) were processed accordingly, followed by a new clustering pipeline together with all SstCre cells, here using the 1000 most differentially expressed genes.

### Measurement of intrinsic properties

Under current-clamp, we applied depolarizing and hyperpolarizing current steps to extract the following electrical properties: resting membrane potential (RMP) was collected after membrane rupture; input resistance (iR) was obtained by the steady-state voltage response to a hyperpolarizing current step injection; H-current-mediated sag was measured as the voltage difference between the peak hyperpolarization and the steady-state response to a long (1 s) current step. Action potential (AP) threshold was obtained from the first AP discharge after the minimum current injection to elicit an AP. The additional following parameters were measured from the same protocol: AP amplitude; AP width at half amplitude; and after-hyperpolarization (AHP) latency (the time from spike threshold to lowest point of the AHP) and amplitude (in mV).

### Statistical Analysis

Data were analyzed for statistical significance using the SPSS (v.17) software package (IBM, Chicago, IL, USA) or Excel. Normal distribution was assumed for data from immunohistochemistry and *in situ* hybridization quantifications, as well as for electrophysiology. Therefore independent t-tests were performed to compare the means of the two groups.

---

**Supplementary Table 1.**
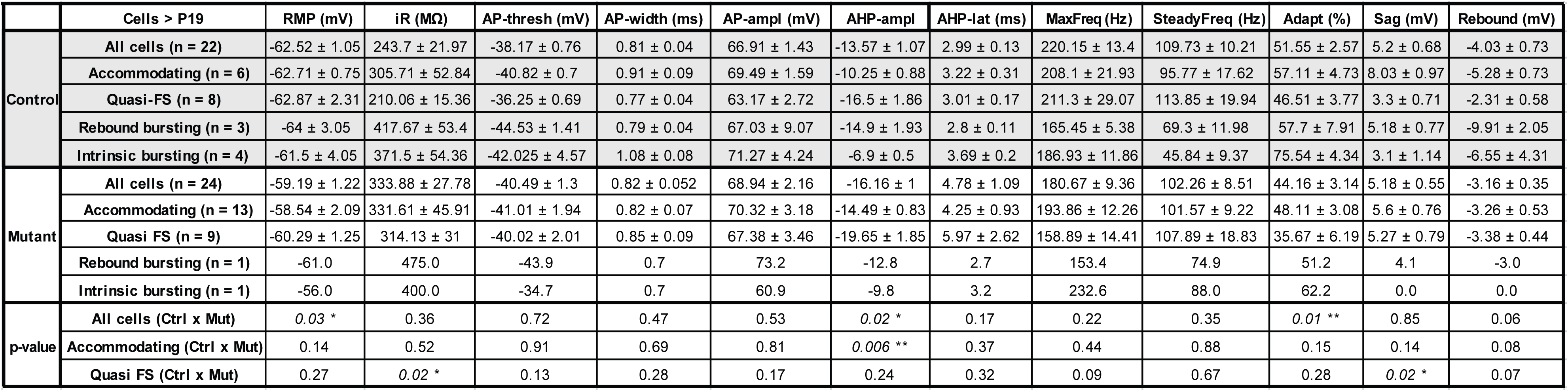
Electrophysiological properties of L5 Sst^+^ neurons. Table shows parameters of L5 eGFP^+^ neurons recorded in S1, in whole-cell patch-clamp mode using Sst^cre/+^;Rosa-Cag-Egfp;Sox6^fl±^ aged P19-30. Values represent mean ± SEM, p-value for unpaired t-tests. The following parameters were analyzed: resting membrane potential (RMP), input resistance (iR), action potential (AP) threshold (AP-thresh), AP half-width (AP-width), AP amplitude (AP-ampl), after-hyperpolarization (AHP) amplitude (AHP-ampl) and AHP latency (AHP-lat), maximum frequency (MaxFreq), steady frequency (SteadyFreq), adaptation (Adapt), sag and rebound. SEM (standard error of the mean).

